# Effects of Serotonin and Dopamine Depletion on Neural Prediction Computations during Social Learning

**DOI:** 10.1101/652693

**Authors:** Anna-Lena Frey, Ciara McCabe

## Abstract

We recently found that individuals with high depression scores demonstrate impaired learning from social outcomes. Given that depression has been linked to altered serotonin (5-HT) and dopamine (DA) functioning, the current study aimed to elucidate the role of these neurotransmitters in social learning with the use of dietary precursor depletion. In a double-blind design, 70 healthy volunteers were randomly allocated to the 5-HT depletion (N=24), DA depletion (N = 24), or placebo (N = 22) group. Participants performed a social learning task during fMRI scanning, as part of which they learned associations between name cues and rewarding (happy faces) or aversive (fearful faces) social outcomes. Behaviourally, 5-HT depleted subjects demonstrated impaired social reward learning compared to placebo controls, with a marginal effect in the same direction in the DA depletion group. On the neural level, computational modelling-based fMRI analyses revealed that 5-HT depletion altered social reward prediction signals in the insula, temporal lobe, and prefrontal cortex. DA depletion affected social reward prediction encoding only in the prefrontal cortex. These results indicate that 5-HT depletion impairs learning from social rewards, on both the behavioural and the neural level, while DA depletion has a less extensive effect. Interestingly, the behavioural and neural responses observed after 5-HT depletion in the current study closely resemble our previous findings in individuals with high depression scores. It may thus be the case that decreased 5-HT levels contribute to social learning deficits in depression.

## Introduction

The ability to learn from social outcomes is crucial for successful interpersonal interactions. We have previously shown that impaired social learning is associated with diminished social engagement motivation and more frequent experiences of negative interpersonal encounters in everyday life [1, 2]. These findings are particularly relevant to the understanding of social impairments in major depressive disorder, as depressed individuals demonstrate reduced learning from social feedback, as well as altered neural encoding of social learning signals [1, 2].

In order to identify potential treatment targets for social learning deficits in depression, it is important to determine which neurotransmitters may contribute to these impairments. Previous research points to a potential involvement of dopamine (DA) or serotonin (5-HT), as these neurotransmitters have been implicated in the psychopathology of depression [3, 4], social processing [5–7], and non-social learning [8–11].

While studies using DA or 5-HT manipulations in combination with *social* learning paradigms are lacking, there is extensive research on the effects of these neurotransmitters on learning from *non-social* outcomes. For instance, behavioural studies have found that lowering DA functioning impairs reward and enhances punishment learning [12–16], whereas increasing DA levels has the opposite effect [17–22]. Moreover, reducing 5-HT functioning has been shown to diminish both reward and punishment learning [23–26], although in some paradigms heightened punishment learning has been observed after 5-HT depletion [27, 28].

On a mechanistic level, it has been suggested that DA and 5-HT neurons contribute to the learning process by propagating learning signals. In particular, it is thought that DA neuron firing represents reward predictions and prediction errors (PEs; indicating the discrepancy between predicted and actual rewards), whereas 5-HT neuron firing may encode punishment PEs [10,29,30]. These mechanisms have been formalised by computational models which, in turn, have been utilised to inform fMRI analyses in humans. Using this approach, it has been shown that increased DA levels are associated with enhanced reward prediction representations in the ventromedial prefrontal cortex (PFC), as well as with heightened reward PE signals in the striatum [17,19,31]. By contrasts, reducing DA functioning has been found to diminish prediction responses in the caudate, thalamus, and midbrain, and to attenuate PE encoding in the caudate, thalamus, and amygdala [13, 32].

In addition, lowering 5-HT levels has been reported to decrease reward prediction representations in the dorsolateral and ventromedial PFC, anterior cingulate cortex (ACC), insula and precuneus [25, 32], while also diminishing punishment prediction encoding in the orbitofrontal cortex and amygdala [33]. Moreover, reduced 5-HT functioning has been associated with attenuated reward PE encoding in ACC, putamen and hippocampus [25, 34].

The above findings demonstrate that DA and 5-HT are involved in behavioural and neural learning processes when *non-social* outcomes are involved. However, it is less clear what role these neurotransmitters play during *social* learning. The current study aimed to examine this question by lowering DA or 5-HT levels in healthy volunteers through acute tyrosine/ phenylalanine or tryptophan depletion, respectively. After consumption of the depletion drink (or a placebo), participants performed a social learning task in the MRI scanner during which they learned associations between name cues and rewarding (happy faces) or aversive (fearful faces) social outcomes. Computational modelling was applied to the data to assess depletion effects on the neural encoding of social learning signals. It was hypothesised that both depletion manipulations would impair social reward learning, while social aversion learning may be enhanced after DA depletion and reduced after 5-HT depletion.

## Methods

### Participants

Seventy right-handed, healthy individuals between the age of 18 and 45 years took part in the current study. Volunteers were screened with the structured clinical interview for DSM-IV (SCID; [35]), and answered several questions about their medical history. Subjects were ineligible if they had a history of any DSM Axis I disorder, a significant current or past medical condition, or any contraindications to MRI scanning. Further exclusion criteria were the current use of any medications besides contraceptives, the use of any psychotropic medications or recreational drugs within the past three months, or smoking more than five cigarettes per week.

In a double-blind, between-subject design, eligible participants were randomly allocated to the DA depletion (N=24), 5-HT depletion (N=24), or placebo (N = 22) group.

The study was approved by the University of Reading Ethics Committee (UREC 15/61) and all subjects provided written informed consent.

### Amino Acid Depletion Drink

The *relative* amino acid amounts for the depletion drinks were based on previous 5-HT [36] and DA [37] depletion studies. However, to reduce the experience of side effects, the *absolute* amounts were adjusted to each participant’s body weight (which has been shown to lead to a reliable depletion effect with a slightly different mixture; see [38]).

Specifically, the placebo drink contained the following amounts for a subject weighing 83.6kg (i.e. the average male weight in the UK), which were adjusted proportionally for lower or higher body weights: L-alanine, 4.1 g; L-arginine, 3.7 g; L-cystine, 2.0 g; glycine, 2.4 g; L-histidine, 2.4 g; L-isoleucine, 6 g; L-leucine, 10.1 g; L-lysine, 6.7 g; L-methionine, 2.3 g; L-proline, 9.2 g; L-phenylalanine, 4.3 g; L-serine, 5.2 g; and L-valine, 6.7 g; L-threonine, 4.9 g; L-tyrosine, 5.2 g; L-tryptophan; 3.0 g.

The 5-HT and DA depletion mixtures were identical to that of the placebo drink, except that they did not contain tryptophan or tyrosine and phenylalanine, respectively. All drinks were prepared by stirring the amino acids and a pinch of salt (to neutralise the bitter taste) into 120mL of tap water, 30mL of caramel syrup, and a tablespoon of oil (with liquid quantities being adjusted to the amino acid amounts).

### General Procedure

After an initial screening visit, eligible participants were sent online versions of the Beck Depression Inventory (BDI; [39]) and a demographics form to complete at home. Subjects were then invited to attend the testing session. They were asked not to consume any food or drinks besides water after 10pm on the previous day, and to arrive at the study location at 9am on the testing day. At this point, participants completed the Positive and Negative Affect Scale (PANAS; [40]) and gave a blood sample which was used to assess baseline amino acid levels. Subsequently, subjects consumed one of the three depletion drinks and were given a protein free breakfast bar. During the following 3.5 hours, participants occupied themselves in a waiting room, with lunch (protein free pasta and tomato sauce) provided at 12 noon. This waiting period was chosen to ensure that the MRI scan took place 5 hours after the consumption of the depletion drink, which is when the maximum depletion effect has been shown to occur [41].

After the waiting period, subjects filled in the PANAS and a side effects questionnaire. Subsequently, they completed a name learning test (see supplement) and the practice trials of the social learning task. Additionally, a second blood sample was collected which was used to assess whether relevant amino acid levels had been successfully depleted (see supplement). Participants then performed the experimental trials of the social learning task in the MRI scanner, and, after the scan, completed a task feedback and drink guess questionnaire (see Figure 1).

**Figure 1:**
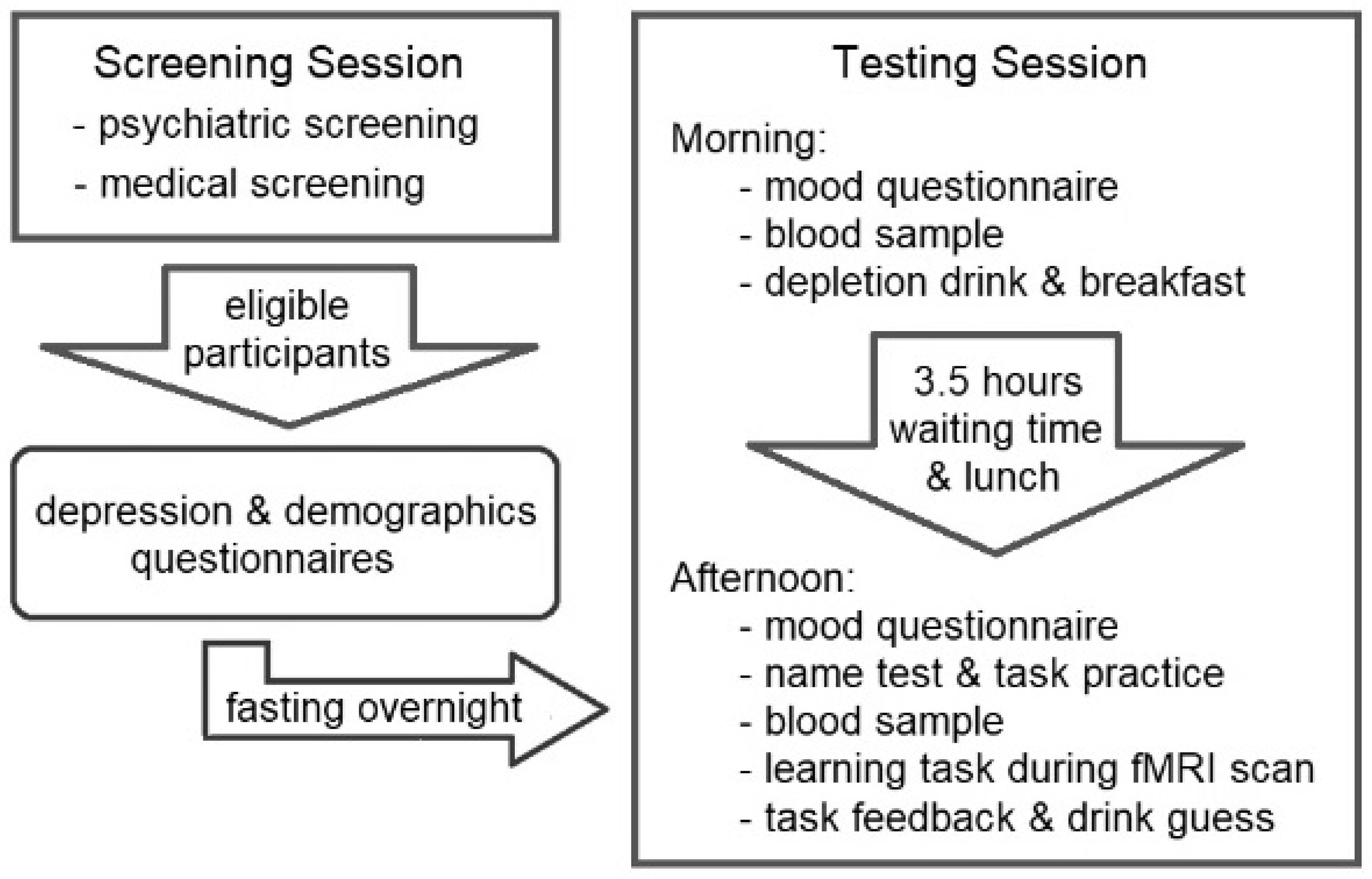
Flow chart of the study procedure; see text for details.

### Social Learning Task

Participants’ aim during the task was to learn associations between name cues and happy, neutral or fearful facial expression. The task consisted of 48 practice and 72 experimental trials, which were divided into social reward and aversion blocks. The blocks were performed in counterbalanced order and three name - face (identity) pairings were randomly allocated to each block. On each trial, participants were presented with a name cue and a rating scale (see below), followed by the face that had been paired with the name (see Figure 2). In the social reward block, each face had a different likelihood (25%, 50% or 75%) of displaying a happy rather than a neutral expression. Similarly, in the social aversion block, each face had a different likelihood (25%, 50% or 75%) of showing a fearful rather than a neutral expression. Participants were asked to learn how likely it was that a given name was associated with an emotional (rather than a neutral) expression and to indicate this likelihood on a visual analogue scale (ranging from 0% to 100%) on each trial before being shown the face. Subjects were instructed to start with a guess and to subsequently base their ratings on the intuition they gained from all the times they had seen the name - face pairing before.

**Figure 2:**
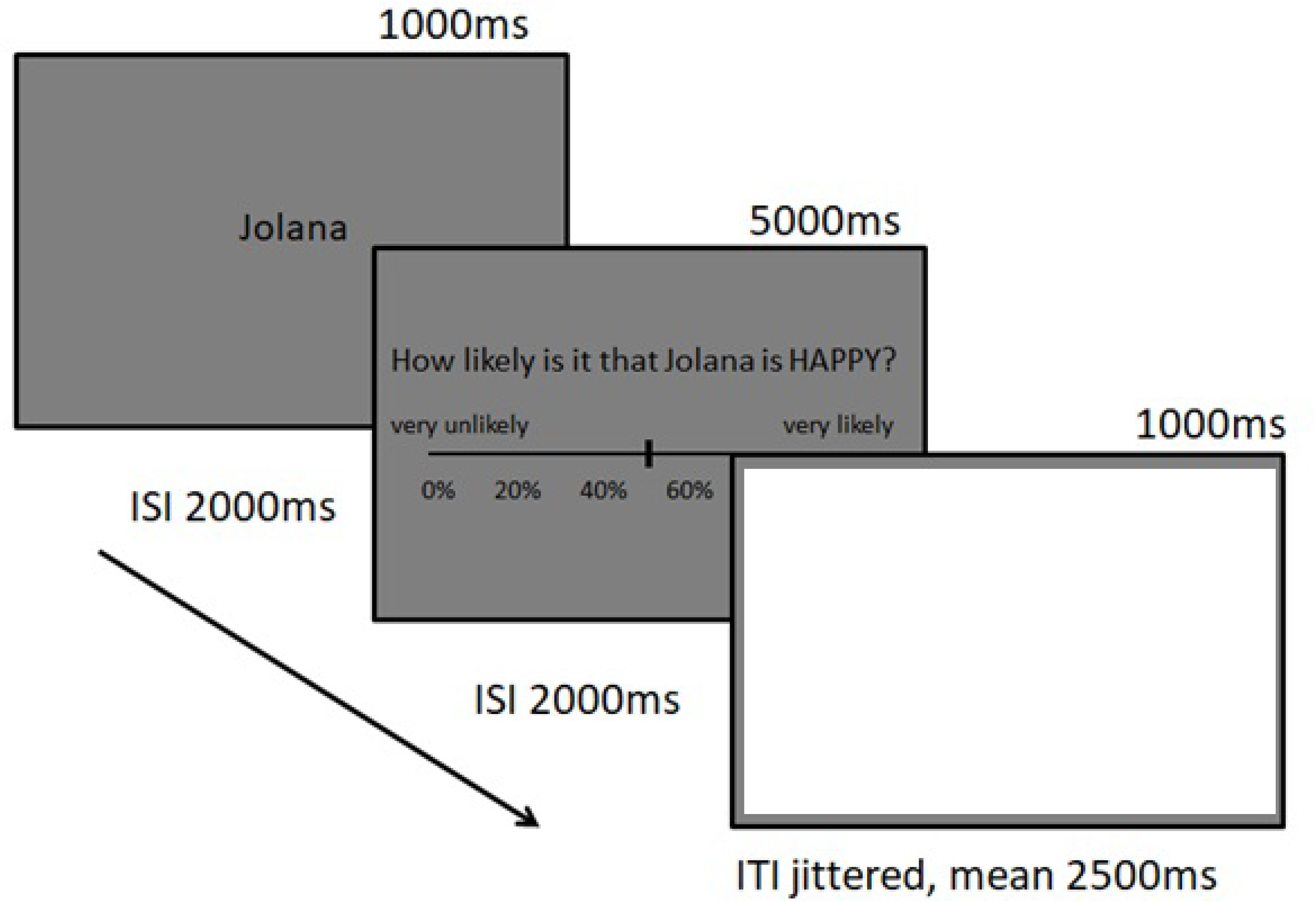
Example of a social learning task trial; see text for details.

### Analysis

#### Behavioural Analysis

Where normality assumptions were met, questionnaire and task measures were analysed using one-way ANOVAs. Otherwise Kruskal-Wallis H tests were used. Additionally, chi-square tests were performed on categorical data.

Visual inspection of the learning task likelihood ratings revealed several clear outliers. Therefore, values outside +/- 2 standard deviations of the mean were removed from the data (removed: N_5-HT depletion_ = 3, N_placebo_ = 3, N_DA depletion_ = 4). Subsequently, a group x valence x probability mixed-measure ANOVA was conducted, and interactions were followed up with one-way ANOVAs. As the sphericity assumption was violated for the probability factor, Greenhouse-Geisser corrected results are reported for the associated effects.

#### Computational Modelling

A Rescorla-Wagner model [42] was fit to the data by minimising the sum of squared errors between participants’ likelihood ratings and the model prediction value (multiplied by 100; similar to [33]). The model included a learning rate (α) and a decay (ƴ) parameter, the latter of which accounted for potential forgetting of the contingencies between the practice and experimental trials (see supplement for details). Group differences in the model fit and parameters were assessed using Kruskal-Wallis H tests.

#### fMRI Acquisition and Analysis

Functional MRI images were acquired using a three-Tesla Siemens scanner (Siemens AG, Erlangen, Germany) and analysed with Statistical Parametric Mapping software (SPM12; http://www.fil.ion.ucl.ac.uk/spm; see supplement for details).

Neural prediction encoding was assessed by entering computational modelling derived prediction values into the first-level fMRI analysis as parametric modulators at the time of the cue (separately for social reward and aversion blocks). On the second level, whole-brain one-way ANOVAs were performed to assess group effects (placebo vs. DA depletion, placebo vs. 5-HT depletion, and DA vs. 5-HT depletion). Reported results were thresholded at 0.005 (uncorrected) on the voxel level and are family wise error corrected at the cluster level.

Additionally, to examine prediction error (PE) encoding, the two PE components (i.e. inverse predictions and outcome values) were used as parametric modulators at the time of the face presentation in the first-level analysis (separately for social reward and aversion blocks). Subsequently, MarsBar (Brett, Jean-Luc, Valabregue, & Poline, 2002) was used to extract average parameter estimates for the two components from a 6mm sphere around striatal coordinates that have been found to encode PEs in a previous meta-analysis (left ROI: −10 8 −6; right ROI: 10 8 −10; Chase et al., 2015). The extracted values were then compared between groups by conducting one-way ANOVAs.

## Results

### Behavioural Results

#### Questionnaires and Demographic Measures

No significant group differences were observed for age (*H*(2) = 0.11, *p =* 0.949), BDI scores (*H*(2) = 1.11, *p =* 0.574), or for the change in pre- to post-depletion PANAS ratings on the positive (*F*(2, 66) = 1.38, *p* = 0.260) or negative (*F*(2, 66) = 0.57, *p =* 0.567) affect subscale (see Table 1). Moreover, chi-square tests demonstrated no significant relations between the depletion groups and sex (*χ*^2^(2) = 0.41, *p* = 0.814), drink guess (*χ*^2^(2) = 0.86, *p* = 0.071), or side effect reporting (*χ*^2^(2) = 1.56, *p* = 0.458).

**Table 1:** Questionnaire and demographic measures by group.

|  | 5-HT Depletion<br>(N = 24) |  | Placebo<br>(N = 22) |  | DA depletion<br>(N = 24) |  |
| --- | --- | --- | --- | --- | --- | --- |
|  | *Mean* | *SD* | *Mean* | *SD* | *Mean* | *SD* |
| N female/ male | 19/ 5 | - | 18/ 4 | - | 19/ 5 | - |
| N reported side effects | 2 | - | 4 | - | 5 | - |
| N drink guessed correctly | 6 | - | 8 | - | 6 | - |
| Age (years) | 21.50 | 3.52 | 21.95 | 4.18 | 21.70 | 4.53 |
| BDI | 2.13 | 2.29 | 2.32 | 2.59 | 3.26 | 3.84 |
| PANAS difference - pos | -2.75 | 5.23 | -1.23 | 4.82 | -3.65 | 4.77 |
| PANAS difference - neg | -1.63 | 3.10 | -0.77 | 2.35 | -1.09 | 2.68 |

SD, standard deviation; BDI, Beck Depression Inventory; PANAS difference - pos/neg, difference between pre- and post-depletion ratings on the positive and negative subscales of the Positive and Negative Affect Scale

#### Social Learning Task Performance

As expected, the mixed-measure ANOVA (group x valence x probability) of participants’ likelihood ratings revealed a significant main effect of probability (*F*(1.36, 77.65) = 209.71, *p <* 0.001), as participants made higher likelihood ratings when the probability of an emotional outcome was greater. Additionally, significant valence by probability (*F*(1.92, 109.45) = 3.35, *p =* 0.040), group by probability (*F*(2.73, 77.65) = 4.42, *p =* 0.008), and group by valence by probability (*F*(3.84, 109.45) = 3.72, *p* = 0.008) interactions were observed.

Follow-up one-way ANOVAs showed significant group differences in the 75% (*F*(2, 57) = 4.81, *p* = 0.012), 50% (*F*(2, 57) = 3.29, *p* = 0.044) and 25% (*F*(2, 57) = 7.03, *p* = 0.002) social reward conditions, with no group effect in any of the social aversion conditions (all F < 2.65). Bonferroni corrected post-hoc tests indicated that, compared to placebo, 5-HT depleted subjects made significantly *lower* likelihood ratings on trials with a 75% chance of displaying a happy expression (*p* = 0.010), but made significantly *higher* ratings on trials with a 25% chance of presenting a happy face (*p* = 0.002). Moreover, DA depleted participants made significantly higher ratings than placebo controls on trials with a 25% chance of displaying a happy face (*p* = 0.040), as well as significantly higher ratings than 5-HT depleted individuals on trials with a 50% chance of presenting a happy expression (*p* = 0.045). These findings indicate that the depletion manipulation, especially 5-HT depletion, impaired social reward learning, seemingly leading to increased *uncertainty* about what social outcomes to expect (as indicated by ratings close to 50% across all outcome probabilities; see Figure 3 below and uncertainty score analysis in the supplement).

**Figure 3:**
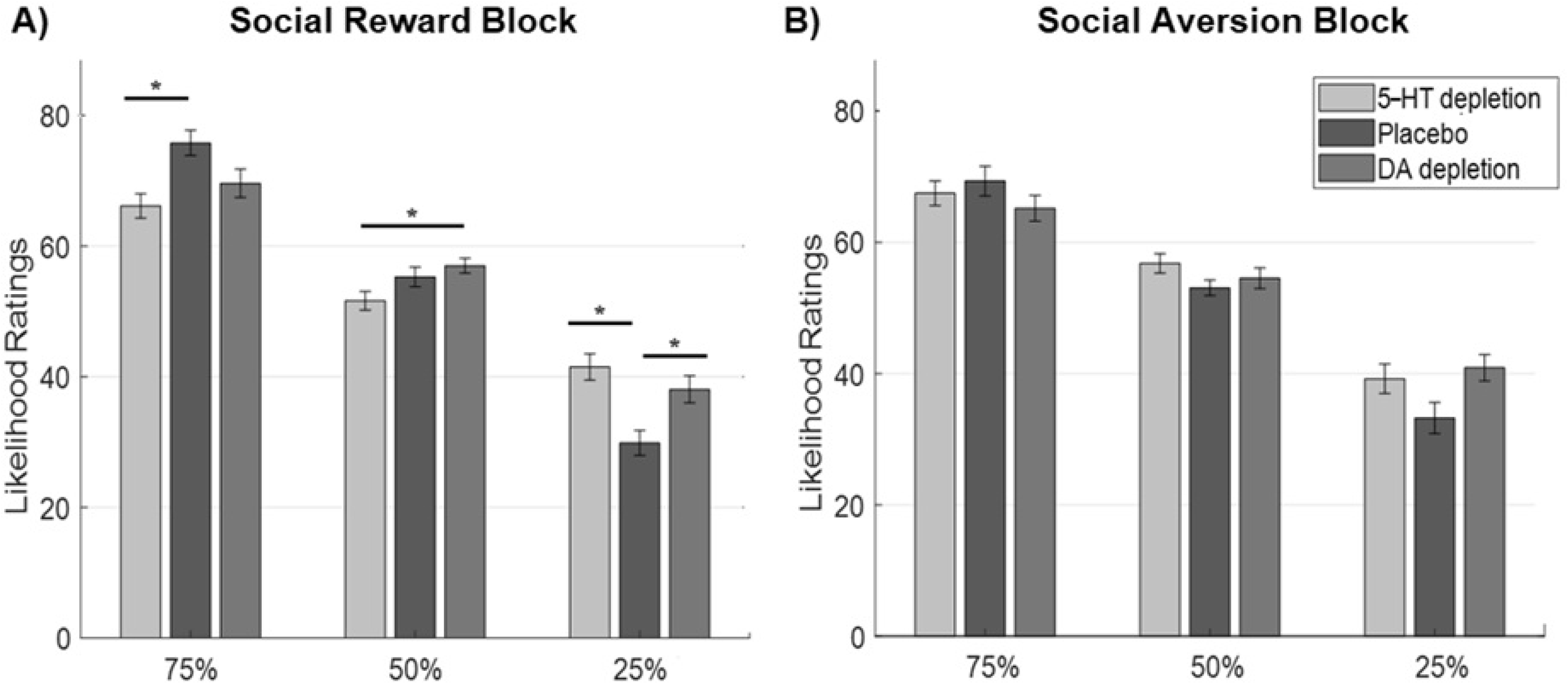
Likelihood ratings by group and probability in A) the social reward block and B) the social aversion block.

#### Computational Modelling

There were no significant group differences in the learning rate (social reward block: *H(2) =* 1.89, *p* = 0.389; social aversion block: *H(2) =* 0.80, *p* = 0.672), or decay (reward block: *H(2) =* 3.37, *p* = 0.185; aversion block: *H(2) =* 1.56, *p* = 0.459) parameters. Similarly, no significant group effects were observed for the model fit, as indicated by mean squared errors, when using individual (reward block: *H(2) =* 2.77, *p* = 0.250; aversion block: *H(2) =* 1.14, *p* = 0.565) or averaged (reward block: *H(2) =* 2.35, *p* = 0.309; aversion block: *H(2) =* 1.81; *p* = 0.406) parameters.

### fMRI Results

#### Neural Prediction Value Encoding

Compared to placebo controls, 5-HT depleted subjects displayed significantly decreased social reward prediction encoding in the dorsal anterior cingulate cortex (ACC)/ dorsomedial prefrontal cortex (PFC), premotor cortex/ dorsolateral PFC, bilateral temporal lobe/ fusiform gyrus, and in the right insula. Moreover, DA depleted individuals demonstrated significantly reduced social reward prediction representations in the dorsal ACC and dorsomedial PFC/ pre-supplementary motor area compared to controls (see Figure 4 below and Table S1 in the supplement). Contrasts between the depletion groups did not reveal any significant clusters.

**Figure 4:**
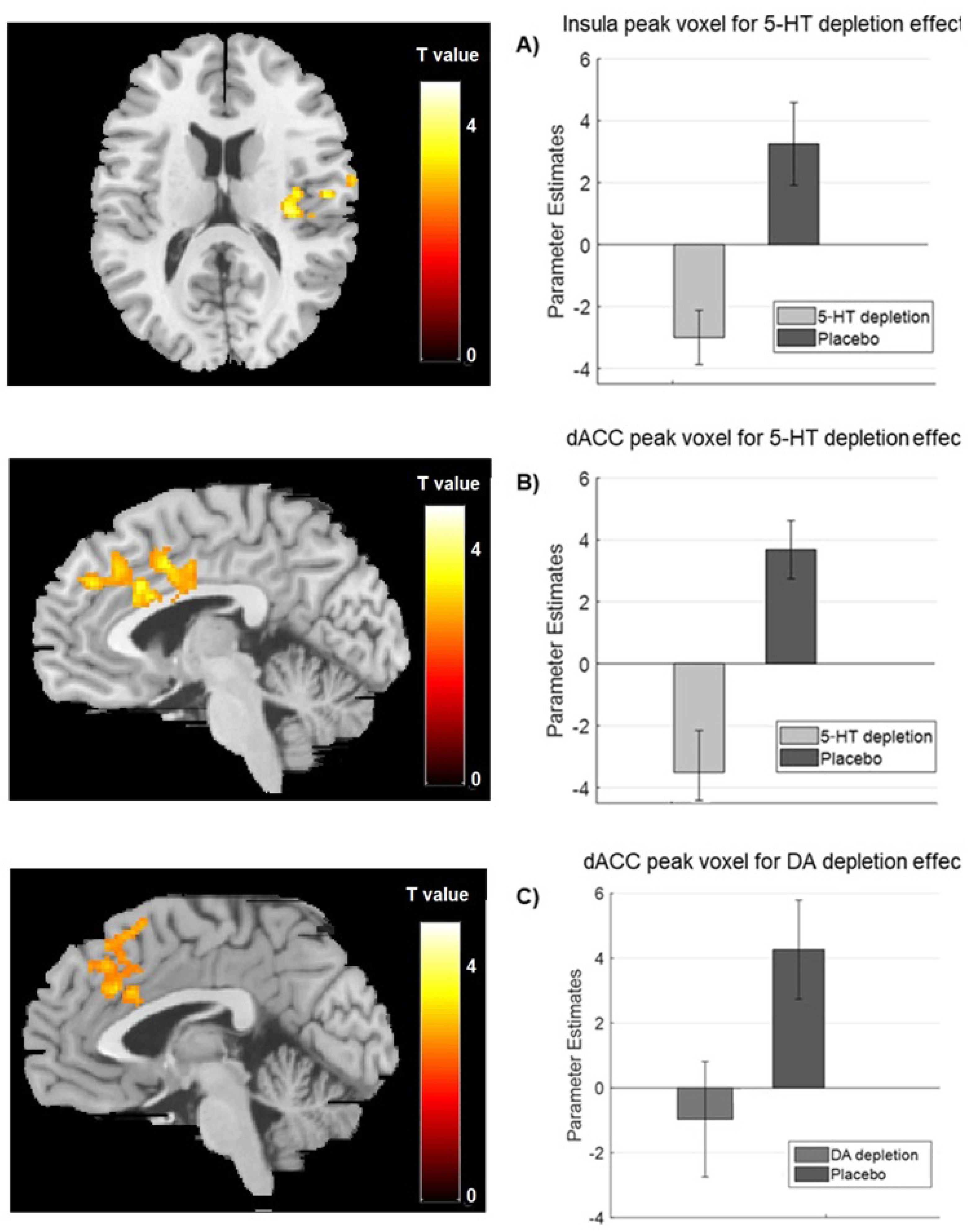
Clusters showing lower social reward prediction encoding in 5-HT depleted (A & B) or DA depleted (C) subjects than in placebo controls, as well as parameter estimates extracted from the peak voxel of the group contrasts in the insula (A) and the dorsal anterior cingulate cortex (dACC; B & C).

Additionally, in the social aversion condition, 5-HT depleted participants demonstrated stronger prediction encoding than placebo controls and DA depleted individuals in the thalamus and precentral gyrus, respectively (see Table S1 in the supplement). All other contrasts yielded no significant clusters.

#### Neural Prediction Error Encoding

One-way ANOVAs were conducted on the average parameter estimates extracted from the striatal regions of interest for the encoding of outcome and inverse prediction values (i.e. the two prediction error components). This analysis revealed no significant group differences for either the social reward or the social aversion block (all F < 0.8).

## Discussion

### Effects of 5-HT Depletion on Social Learning

The present study aimed to examine the effects of 5-HT and DA depletion on learning from social outcomes. The behavioural findings revealed that 5-HT depletion impaired participants’ ability to learn from social rewards, giving rise to heightened uncertainty about what social outcomes to expect. These results are in line with previous reports of decreased non-social learning after reductions in 5-HT functioning [23–26]. Interestingly, using the same task, we previously observed very similar results in individuals with high depression scores [2], which suggests that low levels of 5-HT may contribute to social learning deficits in depression.

Moreover, consistent with the behavioural findings, 5-HT depletion also affected neural learning signals. Specifically, 5-HT depleted subjects demonstrated altered social reward prediction encoding in the dorsal ACC, PFC, insula, and temporal lobe. These observations are in keeping with previous reports of reduced reward prediction signals in the ACC, PFC, and insula following lowered 5-HT functioning [25,32,34].

The engagement of the insula and temporal lobe during the prediction phase of our task may have been due to the role of these regions in the working memory maintenance of faces [43], which may have aided the learning process. Moreover, the dorsal ACC may have contributed to cue value computations [44, 45], while the dorsolateral PFC may have directed attentional resources toward cues that were particularly salient due to their association with happy faces [46].

At first sight, this may suggest that the altered prediction encoding in 5-HT depleted subjects in the above regions may be linked to reduced attentional and working memory processing. However, it should be noted that 5-HT depletion did not merely lower, but instead *reversed*, the neural prediction signals in the abovementioned areas (see supplement). This indicates that, instead of covarying with the prediction of happy faces (as in participants on placebo), brain responses of 5-HT depleted individuals seemed to track the prediction of *neutral* faces.

A possible explanation for this finding is that 5-HT depletion may have given rise to negative biases [47], which may have led to the perception of ambiguous neutral faces as negative. This may have made the latter more salient, resulting in the recruitment of attentional and working memory processes to support the prediction of neural faces. Interestingly, using the same task, we previously found a similar pattern of reversed social reward prediction encoding in the insula and temporal lobe of individuals with high depression scores [2]. Taken together, these findings suggest that low levels of 5-HT may contribute to impaired social reward learning in depression by biasing learning towards negatively perceived ambiguous stimuli.

Following on from the above interpretation, it may seem surprising that no group differences were found in the happy vs. neutral face contrast. However, it is possible that the increased engagement of the PFC in anticipation of neutral faces may have facilitated a preparatory downregulation of limbic regions in 5-HT depleted subjects. This preparatory response may have equalised the otherwise potentially stronger activation to neutral faces in 5-HT depleted subjects compared to placebo controls.

### Effects of DA Depletion on Social Learning

The current study further found that DA depleted participants tended to be less certain about what social rewards to expect compared to placebo controls. This observation is in line with previous findings showing that decreased DA levels are associated with impaired learning from non-social rewards [12–16], while increased DA functioning enhances learning from positive outcomes [17–20,22,31].

Moreover, on the neural level, DA depletion reduced social reward prediction encoding in the dorsomedial PFC and dorsal ACC. This may have been due to an effect of DA depletion on the stability of frontal prediction representations. More concretely, it is thought that the strength of input representations in the frontal cortex is influenced by the balance between D1 and D2 binding, with low levels of DA inducing preferential D2 (rather than D1) binding, which is associated with weak input representations [48]. Therefore, DA depletion may have impaired the stability of prediction representations in the frontal cortex through a shift to predominant D2 binding. This interpretation is in line with that of Jocham and colleagues [31], who found that the D2 receptor antagonist amisulpride increased predictive value signals in the vmPFC, possibly by facilitating more stable D1- (rather than D2-) mediated value representations.

It should be noted that, in our task, DA depletion had a less extensive effect on behavioural and neural responses than 5-HT depletion. This may suggest either that DA is less crucially involved in *social* learning in particular, or that the stimuli used in our task (happy faces of strangers) were not rewarding enough to elicit a robust DA response. Future studies using different, more unambiguously rewarding social stimuli (such as pictures of friends) are needed to distinguish between these possibilities.

### Conclusion

Taken together, the results of the current study indicate that 5-HT depletion impairs social reward learning on both the behavioural and the neural level, possibly partly by increasing attentional and working memory processing of negatively perceived neutral faces. DA depletion had a similar, although less pervasive, effect. Interestingly, the behavioural and neural responses observed after 5-HT depletion in the current study closely resemble our previous findings in individuals with high depression scores. It may thus be the case that decreased 5-HT levels contribute to social learning deficits in depression. It would be of interest for future studies to examine whether serotonergic antidepressants alleviate social learning impairments in depressed individuals.

## Acknowledgments

We would like to thank Shaama Reese, Elena Dyankova, Zaneta Filipczak and Kirsty Holroyd for assisting with the data collection, Nicholas Michael for running the biochemical amino acid analysis, Rada Mihaylova for providing venepuncture training, and Shan Shen for helping with the MRI scanning.

The current work was funded by the Medical Research Council PhD studentship of Anna-Lena Frey.

